# A theoretical foundation of state-transition cohort models

**DOI:** 10.1101/430173

**Authors:** Rowan Iskandar

**Author notes:** Current Address: Department of Health Services, Policy, and Practice, Brown University, Providence, Rhode Island, USA.

## Abstract

Following its introduction over three decades ago, the cohort model has been used extensively to model population trajectories over time in decision-analytic modeling studies. However, the stochastic process underlying cohort models has not been properly described. In this study, we explicate the stochastic process underlying a cohort model, by carefully formulating the dynamics of populations across health states and assigning probability rules on these dynamics. From this formulation, we explicate a mathematical representation of the system, which is given by the master equation. We solve the master equation by using the probability generation function method to obtain the explicit form of the probability of observing a particular realization of the system at an arbitrary time. The resulting generating function is used to derive the analytical expressions for calculating the mean and the variance of the process. Secondly, we represent the cohort model by a difference equation for the number of individuals across all states. From the difference equation, a continuous-time cohort model is recovered and takes the form of an ordinary differential equation. To show the equivalence between the derived stochastic process and the cohort model, we conduct a numerical exercise. We demonstrate that the population trajectories generated from the formulas match those from the cohort model simulation. In summary, the commonly-used cohort model represent the average of a continuous-time stochastic process on a multidimensional integer lattice governed by a master equation. Knowledge of the stochastic process underlying a cohort model provides a theoretical foundation for the modeling method.

## Introduction

Decision models have been used in various applications from clinical decision making to screening guideline development. The objective of decision modeling is to integrate and present evidence within a coherent and explicit mathematical structure that can be used to link evidence to decision-relevant outcomes. In decision modeling, a state-transition Markov model is often used to simulate the prognosis of a patient or a group of patients following an intervention. Beck and Pauker [1] introduced the modeling method over 30 years ago with the aim to provide a simple tool for prognostic modeling and for practical use in medical decision making. They applied standard methods from the Markov chain theory [2, 3] to simulate a life history of *an individual* which is structured as transitions across various heath states over time, i.e. a stochastic process on a finite state space (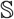). However, the current literature in decision modeling lacks clarity in the following two concepts. First, a *cohort model*, an extension of the state-transition model from one individual to a group of individuals, is often used to introduce and describe a stochastic process on 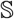. According to [1] and the subsequent published tutorials and textbook [4–7], given a matrix of transition probabilities and an initial distribution of *counts* of individuals across health states, a cohort model generates the life trajectory of a cohort of *identical* individuals by repeated multiplications of the vector of population counts by the transition probability matrix. This matrix operation alludes to a stochastic process on a much larger state space, i.e. 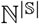 (compared to 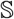),where ℕ and 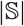 denote the set of natural numbers and the number of states, respectively. For example, 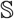 may be a partitioning on an individual’s health state into healthy, sick, or dead. In contrast, 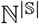 may refer to a partitioning of a cohort of individuals into the numbers of healthy, sick and, dead individuals. The convolution of these two different processes in decision modeling literature lead to the second issue: practitioners are taught that cohort models capture the *average behavior* of the individuals. [7, 8] However, this claim is often stated without any clear reference to which stochastic process. Frederix et. al. [9] proposes an ordinary differential equation (ODE)-based method to approximate Markov models whilst acknowledging that the ODEs can describe only “the typical (mean) change,” albeit, without providing any rational or reference. One of the standard textbooks [7] describes cohort model as a representation of the average experience of patients in the cohort without providing any rationales to support this description. Furthermore, wide acceptance of methodologies does not automatically imply veracity. [10] In the context of cohort models, the wide acceptance is ingrained by the ISPOR-SMDM Modeling Good Research Practices Task Force-3. [11] The published best practices cites the work of Beck et al. [1, 4] as the main and *only* references for cohort models, thereby standardizing cohort models as the recommended method for simulating population trajectories despite of the lack of a proper description of the theoretical support in the decision modeling area.

This paper aims to address the aforementioned problems by placing cohort models in a larger framework based on existing literature on stochastic process. Therefore, in this paper, we explicate the stochastic process underlying a cohort model and derive its corresponding probability function from which the prevailing notion of an *average* is based on. For this paper, we adopt a continuous-time perspective, i.e. for a discrete-time process, a corresponding continuous-time process exists (*embeddability assumption*). In addition, we assume that transition rates are time-invariant and the process of interest is Markovian, i.e. knowledge of the current state conveys all information that is relevant to forecasting future states. This paper is organized into four main parts. Firstly, we review the prevailing descriptions of cohort model in decision modeling literature and formulate their mathematical representations to substantiate our claim of lack of clarity in the current description of a cohort model. Secondly, we postulate a precise description of the cohort dynamics and the probability rules on these dynamics to derive the underlying stochastic process. Thirdly, we solve for the probability function associated with the stochastic process by method of evolution equation. To show that the derived stochastic process corresponds to cohort model, we also derive the mathematical formulas for the average and the variance of the process. Fourthly, the correspondence is then demonstrated by showing the equivalence between the first two moments of the process and the cohort model via a numerical example. Throughout this exposition, we try to strike a balance between mathematical rigor and accessibility to practitioners.

## Methods

### Prevailing views of cohort model

Beck and Pauker [1] introduced a discrete-time Markov model as a method for representing a patient’s prognosis which can be viewed as a sequence of states of health in discrete time steps. We distinguish between three prevalent descriptions of a Markov model [4, 5, 7, 8]: (1) a model for an individual or a Markov chain on 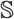, (2) a cohort model or a process on 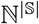, and (3) the continuous-time analogue of a cohort model from which the ODE-based method [9] arises.

### Markov chain on 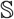

A discrete-time Markov chain on a finite set of mutually exclusive and completely exhaustive states: 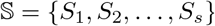, is a stochastic process, {*S*(*n*)} = {*S*(*t_n_*)} where 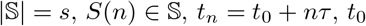 is the initial time, *τ* is the Markov cycle and *n* ∈ {0, 1, …, *N*}, given an *s* × *s* stochastic matrix of transition probabilities governing the transitions among states, **P**_*τ*_(*t*) = (*p_ij_*(*t*))_1≤*i,j*≤*s*_, and a 1 × *s* vector of initial probability distribution **p**(*t*_0_) = [1 0 … 0 0] if, without loss of generality, we assume an individual start in *S*_1_. Each *p_ij_*(*t, τ*) has the usual interpretation of the probability of an individual transitioning from *S_i_* to *S_j_* in *τ* at time *t*. In principle, to calculate the distribution of states in the next cycle, **p**(*t*+*τ*), we post-multiply the distribution in the current cycle, **p**(*t*), with **P**_*τ*_(*t*):

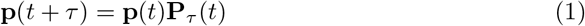

### Cohort simulation

An *s*-state cohort simulation [1, 7] is a recursive algebraic calculations that generates a trajectory of population counts (*state-configuration*): 
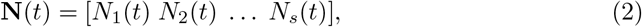

where **N**(*t*) ∈ ℝ^*s*^ denotes the number of individuals in state *S_i_* at time *t* ≥ *t*_0_, given **P**_*τ*_(*t*) as defined above, and an initial distribution **N**(*t*_0_) = [*n*_0_ 0 … 0 0]. In principle, to calculate the distribution of individuals across all states at time *t* + *τ, N*(*t* + *τ*), the state-configuration at time *t, N*(*t*), is projected forward in time using the linear operator **P**_*τ*_(*t*): 
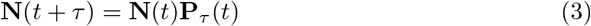

### Continuous-time ODE-equivalence

The recursive formula (Eq 3) can be written as a difference equation and if we take the limit of this difference equation with respect to the Markov cycle, we obtain the differential form of a cohort model: 
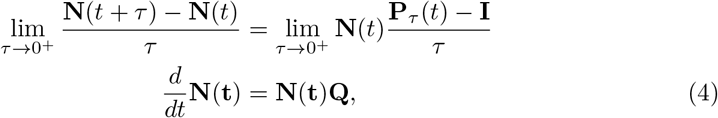

where **Q** is the *infinitesimal generator* matrix of size *s* × *s*, **Q** = (*ĉ_kl_*)_1≤*i,j*≤*s*_, for the continuous-time stochastic process underlying a cohort model. The off-diagonal elements of **Q** are non-negative while the diagonal elements are non-positive, 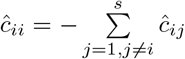. If 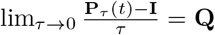 [12, 13], then the continuous-time version of a cohort model is a *deterministic* process governed by an ordinary differential equation (ODE) or, alternatively, a rate equation for the state-configuration.

## Reformulation of cohort model

### Formulation strategy

In the context of decision modeling, the typical system of interest is exemplified by the health state of an individual following the implementation of an intervention and the changes in the health state during a time period, often a lifetime. The representation is dynamic in the sense that the states and parameters of the system are functions of time. The formulation of a *dynamic* process underlying the changes in an individual’s health state starts by observing the state of the world at, at least two different time points, e.g., at *t* and *t* + *τ*. The difference between the two observations are then identified and the events, that contributed to these differences in health state and are most consistent with the prevailing evidence and theories, are postulated. After identifying all possible events, an individual’s health state is then represented by enumerating the possible health states, i.e. a partitioning on the world via S. The dynamics of the system is consequently determined by the identified events and operationalized by assigning the allowable transitions to all pairs of states in 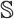 that are consistent with theories and data. The aforementioned formulation of a process for an individual, which follows Beck and Pauker [1], is then extended to a cohort of individuals, i.e. a process on ℕ^*s*^. In this expanded state space, the dynamic system of interest is the changes in state-configuration, which is represented by Eq 2. The transition from one state-configuration to another represents not only a change in health state of one individual, which follows the rules on 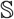 but also a change in the number of individuals of the two corresponding states.

Given the state space and the transition rules, we then specify the nature of the transitions, i.e. stochastic or deterministic changes. The choice of a stochastic (vs. deterministic) representation of the transition is natural because the applications of decision modeling often involves a disease progression component which is an inherently random system, i.e. can be described up to a certain degree of confidence. In contrast to a deterministic model, in which given an initial condition, the sequence of the states of the world can be determined with a probability of one, the solution to a stochastic model can be represented by a multidimensional probability function at any *t* following the initial time *t*_0_. Therefore, the state of the system at any given time can only be described with a probabilistic statement, i.e., the probability of observing a particular state-configuration, **N**(*t*) = **n** ∈ ℤ^*s*^ at time *t* or *P*(**N**(*t*) = **n**) = *P* (**n**, *t*).

### Master equation

Given the formulation strategy, we formalize the description of the process in this section. An event of transitioning from *S_k_* to *S_l_* is denoted by the *transition R_kl_* where *k, l* ∈ {1 …, *s*}. Since the occurrence of each transition alters the state-configuration, each *R_kl_* is associated with a vector representing this change, *s_kl_*, and is defined as *s_kl_* = *e_l_ − e_k_* where *e_i_* is the *i*-th column of an identity matrix of size *s*. We associate each transition with a *propensity function*: *v_kl_*(**n**, *t*) = *c_kl_*(*t*)*h*(*n_k_, n_l_*), where *c_kl_*(*t*) is the transition rate for transition *R_kl_*, and *N_k_*(*t*) = *n_k_* and *N_l_*(*t*) = *n_l_*. The function *h*(*n_k_, n_l_*) determines the number of ways *R_kl_* can occur which is equal to the number of individuals in *S_k_, h_kl_*(*n_k_, n_l_*) = *n_k_*. The propensity function denotes the probability of a (any) transition occurring in continuous time as a function of the state-configuration and rate constant. The probability of observing *one particular* transition *R_kl_* in a small time interval *τ* is assumed to be linear in time: *c_kl_*(*t*)*τ* + *o*(*τ*), where *o*(*τ*) represents other terms that decay to zero faster than *τ*. Therefore, the probability of a transition from *S_k_* to *S_l_* in time step *τ* is given by *n_k_c_kl_*(*t*)*τ* + *o*(*τ*) (see S1 Transition probability). A transition *R_k,l_* occurring with a probability *v_kl_*(**n**, *t*)*τ* in a time step *τ* alters the current state **N**(*t*) = **n** to:

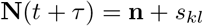

Following the mathematical descriptions of the possible changes in the state-configuration and the probabilities of these changes, we derive the evolution equation for *P* (**n**, *t*), i.e., master equation (also known as the *forward Kolmogorov* equation), which describes how *P* (**n**, *t*) changes in continuous time. The master equation is obtained by enumerating all possible events contributing to a change in the probability of observing a particular state-configuration in an infinitesimal time step and is given by (see S2 Master equation:

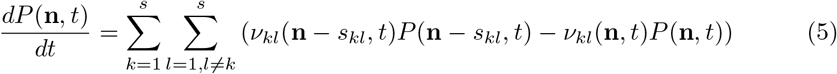

The terms, *v_kl_*(**n**, *t*)*P* (**n**, *t*), represent the probability transferred from one to another state-configuration resulting from the transition *R_kl_*. (*probability flux*). In principle, a master equation expresses the change per unit of time of the probability of observing a state-configuration **n** as the sum of two opposite effects: the probability flux into and out of **n**. Therefore, a master equation is essentially a rate equation for the probability of observing a state-configuration (c.f. Eq 4). The solution to the master equation, *P* (**n**, *t*), contains all information about the modeled system and can be, under some conditions on *c_kl_*(*t*) obtained by applying the generating function technique (see S3 Generating function method). A probability generating function (PGF) of *P* (**n**, *t*) is defined by:

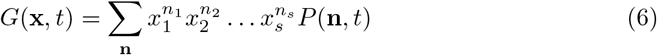

The explicit solution to the Eq 6 is obtained by solving a first-order linear partial differential equation (PDE) for *G*(x, *t*). [14] We solve for the PGF under two initial conditions: (1) an arbitrary initial distribution and (2) all individuals start in *S*_1_. From the solution to the PDE, we recover *P* (n, *t*) by using the definition of PGF and obtain the first two moments of the process, i.e., the mean and the variance from the first derivative of *G*(x, *t*) (see S4 Mean and variance of the master equation).

## Results

### Solution to master equation

The solution to the master equation, the probability function of a state-configuration **n** at time *t*, for an arbitrary initial state-configuration *P* (**n**, 0) is given by: where **z**_*i*_ = [*z*_*i*1_ *z*_*i*2_ … *z_is_*] where *i* ∈ {1, 2, …, *s*}, *B*(*t*) = exp *At*, i.e. the matrix exponential and *A* is an *s* × *s* matrix, (*c_kl_*)_*k,l*∈{1,2,…,*s*}_, with elements of the form: *a_kl_* = *c_kl_* − *γ_k_δ_kl_* with 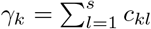 and *δ_kl_* is the Kronecker delta (*δ_kl_* = 1 if *k* = *l* and is 0 otherwise). Matrix *A* is identical to matrix *Q* in Eq 4, the generator matrix for the continuous-time ODE-equivalence of a cohort model. For the case where all *n*_0_ individuals start in *S*_1_ (*N*_1_(0) = *n*_0_), the probability function of a state-configuration **n** at time *t* is represented by: 
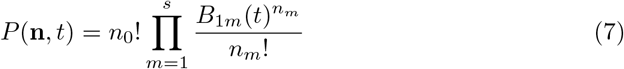

The mean of the number of individuals in the *i*-th state at time *t* is given by: 
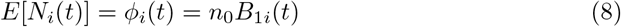

The variance of the number of individuals is the *i*-th state at time *t* is given by: 
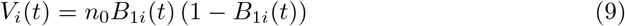

For an arbitrary initial distribution, the mean of the number of individuals in state *S_i_* is given by: 

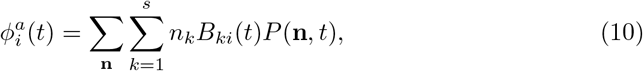
and the variance is given by: 
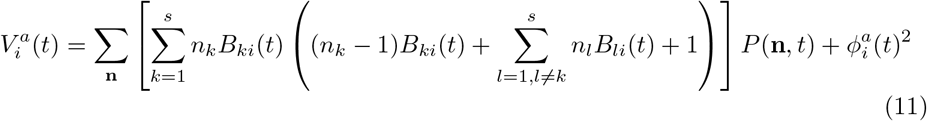

### Numerical verification

We conduct a numerical exercise to verify the validity of the derived stochastic process. In particular, we aim to verify the formulas for the mean (Eq 8) and the variance (Eq 9). For this verification exercise, we consider a stochastic process with *s* = 4, i.e.

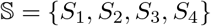 (Fig 1), rate constants (elements of *Q*): *c*_12_ = 0.05, *c*_13_ = 0.01, *c*_14_ = 0.001, *c*_23_ = 0.1, *c*_24_ = 0.05, and *c*_34_ = 2, and an initial condition in which all individuals start in *S*_1_, i.e. *N*_1_(0) = 10000. As a set of benchmarks for the mean, we simulate the population trajectories, i.e. the counts of individuals in all states, (1) using a cohort model with *τ* = 1-year (Eq 3) and (2) a continuous-time cohort model (Eq 4). The benchmark for variance is based on 1000 Monte Carlo samples of *microsimulation* population with *τ* = 1-year. Each microsimulation population is conceived as a realization of 10000 Markov chains with the generator matrix *Q* (as in Eq 4). The calculation of the 1-year transition probability matrix from the *Q* matrix is based on the formulas from Welton and Ades. [15] The results of the population trajectories are given in Figure (Fig 2). We observe no difference between the empirical estimates from the microsimulation and the analytic formulas for both the mean and the variance of the process associated with the cohort model.

**Fig 1.**
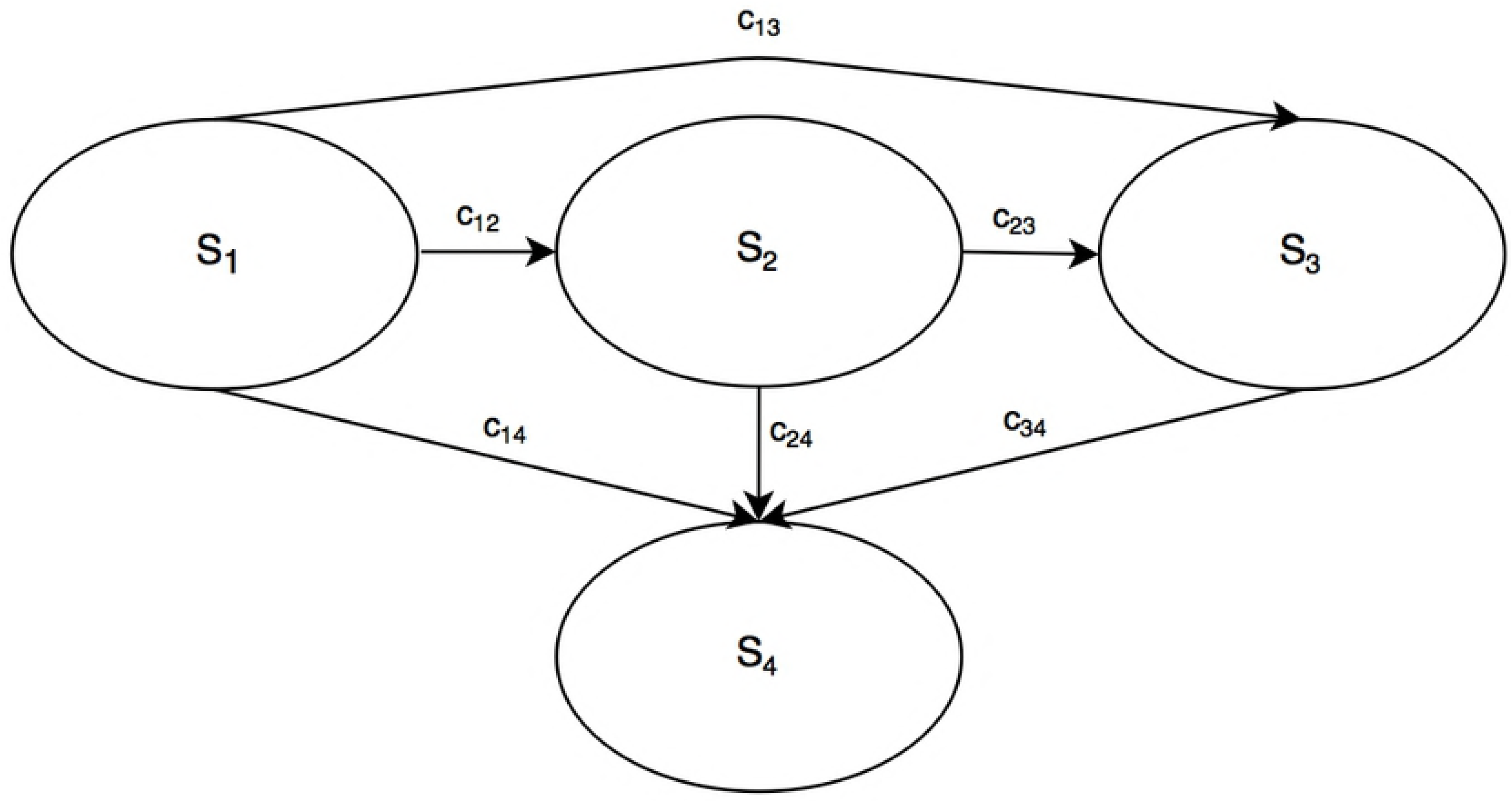
A 4-states continuous-time stochastic process used in numerical verification exercise

**Fig 2.**
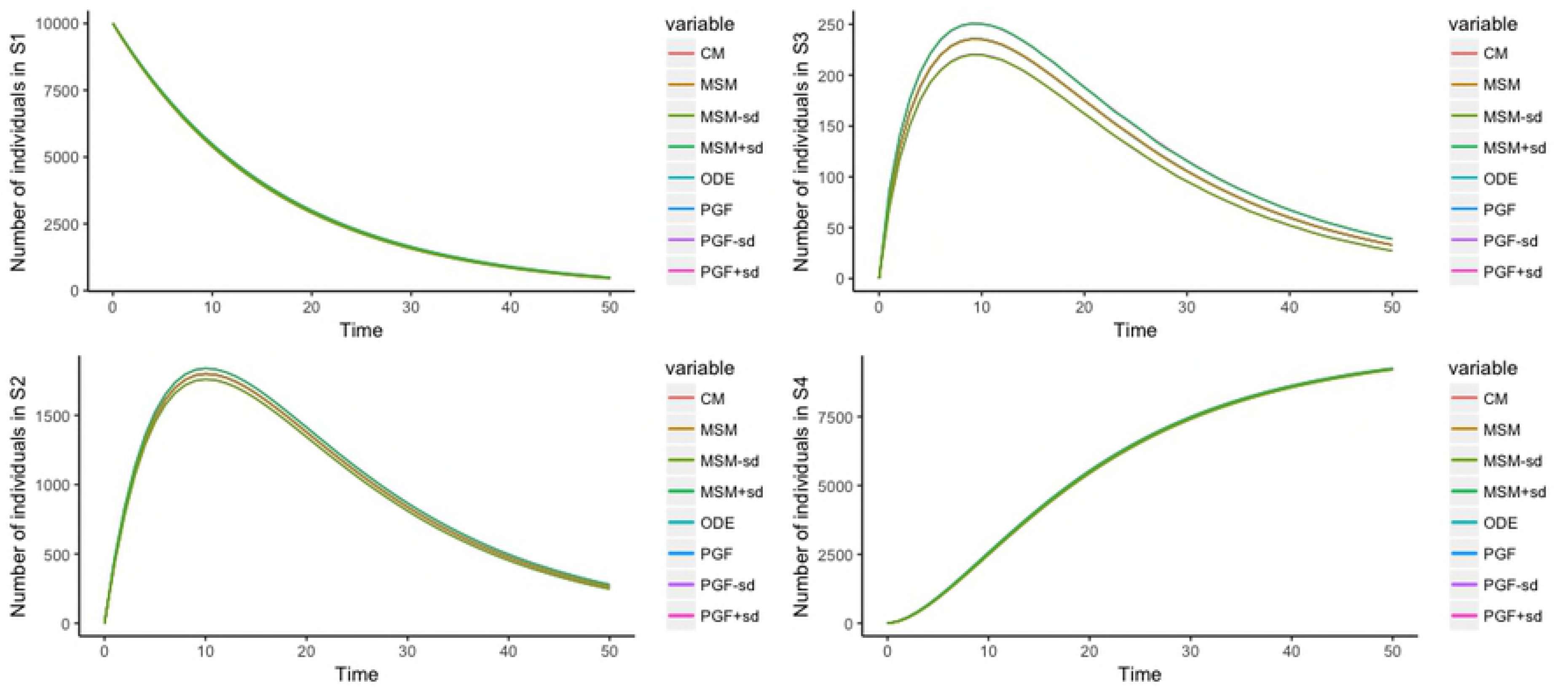
Population trajectories across 4 states [CM: cohort model, MSM: microsimulation, ODE: continuous-time cohort model, PGF: master equation, sd: standard deviation.

## Discussion

In this study, we explicate the stochastic process underlying the commonly used cohort model, defined as a continuous-time discrete-state stochastic process, in the context where population counts are of interest. First, We start the derivation by specifying a stochastic state-transition model for an individual following Beck and Pauker: a stochastic process on state space 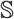. [1] We then extend the model to a cohort of individuals by formulating a set of postulates for the cohort dynamics: a stochastic process on 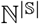. The evolution equation representing the stochastic process for the cohort dynamics in the form of master equation is then obtained. We then solve the master equation to get the probability function of individuals across all states, which completely characterizes stochastic nature of the dynamics, by using the generating function method. Based on the explicit form of the generating function, the analytical formulas for the average and variance are obtained. This measure of variance quantifies the inherent variability of the population trajectories (*aleatory uncertainty* [16] or first-order uncertainty [17]) and may be useful for quantifying the extent of variability in the mean prediction. In our simulation example, we show that to estimate the variance of the population counts across states, we need to replicate the microsimulation population a sufficiently large number of times, which is a computationally intensive task and is similar to applying uncertainty analysis to a microsimulation model. [18] Our theoretical result provides a more direct and non-computationally demanding approach through the explicit formula for the variance. The derived probability function, when all individuals start in one state (a typical scenario in most applications) takes the form of a multinomial distribution and is a known result from the stochastic compartmental model literature [19]. In applications where the relevant population comprises of an actual cohort (e.g. birth cohorts), the individuals are often, prior to an implementation of an intervention, distributed across health states. For this purpose, we also derive the probability function and formulas for the moments, for an arbitrary initial condition. Lastly, we show the equivalence between the average of the derived stochastic process and the cohort model; thereby substantiating the prevailing notion of cohorts models as average process.

In the introduction to cohort models [1] for use in decision modeling, the authors emphasized the clinical utility of a *Markov* process (as opposed to a decision tree model) for modeling patient’s prognosis with ongoing risks: a model for an inherently stochastic system. A stochastic representation of a system amounts to making probabilistic statements about its state given a set of probabilistic rules for the system dynamics. Therefore, the solution to any stochastic processes or models takes the form of a probability function on a set of random variables [2], i.e., in our context, *P* (**n**, *t*), which is the solution to the master equation. However, in their expositions, population counts were used as a representation of the stochastic system via a *cohort simulation*. The multiplication of the transition probabilities (used in individual process) with the cohort size (3) produces a deterministic quantity, i.e. the average of the process, not a probabilistic statement about the process. The commonly used representation of a cohort model via cohort simulation in decision modeling literature convolutes two distinct processes: a process on 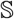 and a Markov chain on ℕ^*s*^. The notion of the average as represented by cohort simulation is based on the latter. To date, our work is the only study that delineates the convolution of these two different processes which has been perpetuated by tutorials and textbooks in decision modeling [7, 20].

Having an explicit form of the evolution equation for *P* (n, *t*) is advantageous for, in some cases (e.g. time-invariant rates), finding the analytical form of *P* (**n**, *t*) and, in most cases (e.g. time-varying rates), deriving the mean and the variance of the process by using their evolution equations and without knowing the explicit form of *P* (**n**, *t*). In addition, given this knowledge of evolution equation associated with a cohort model, we can identify the relationship between cohort models and other known methodologies in stochastic process. In the classification tree of Markov processes [21], a cohort model as represented by the master equation is derived from the differential Chapman-Kolmogorov equation without the drift and diffusion coefficients. If the cohort size is sufficiently large, a cohort model can be approximated by the van Kampen’s expansion method [22], which is analytically more tractable compared to master equation. In addition, if we relaxed the integrality constraint on the state space to allow for nonnegative real numbers, a cohort model may be represented by a Fokker-Planck or a stochastic differential equation (SDE). [21, 23] Therefore, our study results provide practitioners and researchers the proper platform for the porting of methods from stochastic process literature for applications in decision modeling.

Knowledge of the underlying process will provide a proper context for efforts to resolve or clarify methodological challenges or issues, for example, in estimating the appropriate time step for cohort models. [24, 25] Practitioners of cohort models are taught to add an additional half-year of lives at the end of the cohort simulation to account for the fact that individuals transition at the end of each time step. This half-cycle correction is a consequence of discretizing a continuous-time scale. However, the refinement of half-cycle correction method was based on the average process [26], i.e. not taking into account the variance of the process or other higher-order moments. Naimark et al. [26] proposed several methods, including numerical integration techniques to determine the appropriate time step to bypass the need of half-cycle correction. Elbasha and Chhatwal [27] quantified the approximation errors from using these numerical integration methods in the context of a 3-states cohort model. However, these numerical approximation methods were applied to the “state membership function,” which is essentially the average number of individuals: an approximation of an approximation. This study result can facilitate a more accurate approximation, which would be based on directly using the master equation, i.e. based on how a change in the time step affects the probability function.

One limitation of the study concerns with the focus on cohort models with time-invariant rates. In most applications, the transition probabilities are often time-dependent (e.g. age-specific probability of death due to other causes). A direct consequence of assuming non-constant rates is that the solution to a master equation becomes mathematically intractable and henceforth inaccessible to most practitioners. However, various numerical and approximation methods for solving master equations are available as a corollary of knowing the proper placement of cohort models in the classification tree of stochastic processes, as described above. For example, Gillespie’s stochastic simulation [28] and tau-leap methods [29] numerically solves a master equation by simulating all possible events one-at-a-time or in batches, respectively. Despite of this limitation, our analytical results remain applicable in many situations since no restriction is placed on the number of states and the allowable transition patters on the set of states. In addition, the analytical results for an arbitrary initial condition can be utilized to approximate a process with time-varying rates by using a piecewise process. Each piece or time interval corresponds to a time-invariant process. The initial condition of the next time interval would be the distribution of individuals at the end of the previous interval. We plan to introduce this piecewise method in a future study.

This paper also assumes that the discrete-time transition probability matrix is embeddable to a continuous-time model, i.e. the limit of *P* exists and its equal to the generator matrix of the process (Eq 4). Although discrete-time Markov models has been the method of choice in many applications in decision modeling, the processes under consideration, e.g. biological process or disease progression, evolve continuously; henceforth the appropriate modeling framework would be a continuous-time model [30]. In addition, the reason for the greater popularity of the discrete-time structure stems from its simpler mathematical nature, not from considerations of veracity. [31] However, in some instances, one can argue that an individual-level model tend to follow a discrete-time process, e.g. the transitions among living arrangements for an elderly patient occur in discrete time steps. However, if we consider a cohort of individuals, the probability of a patient transitioning in a small time interval would very likely to be nonzero. Nevertheless, we believe that future studies investigating whether the typical Markov models in decision modeling are embeddable are warranted.

## Conclusion

The cohort model provides an easy-to-implement method for modeling recurrent events over time and has been used extensively in many applications ranging from clinical decision making to policy evaluations. Knowledge of the underlying theoretical framework would reaffirm the prevailing beliefs in the veracity and validity of the method and thereby increase confidence in conclusions derived from studies using cohort models.

## Supporting information

**S1 Transition probability** In this section, we show the derivation for the probability of transitioning from *S_k_* to state *S_l_* in time step *τ*, which is given by:

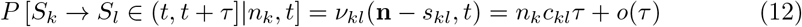

The probability of a *particular* transition *k → l* (*R_kl_*) in a small time step *τ* is postulated to be linear in time: *c_kl_τ* + *o*(*τ*). This postulate is conceptually a first-order Taylor approximation to an instantaneous transition probability. The transition rate constant, *c_kl_*, is interpreted as the derivative of the transition probability at time *τ* = 0. We also restrict the number of allowable transitions in *τ* to one transition. If the number of individuals in *S_k_* at time *t* is equal to *n_k_*, then any of these *n_k_*, assumed to be *indistinguishable*, individuals is at risk of transitioning. Since individuals are indistinguishable, each individual transition can be considered as a Bernoulli trial with a probability of success of *c_kl_τ* + *o*(*τ*). The probability of only one individual transitioning is equal to: (*c_kl_τ* + *o*(*τ*))(1 − *c_kl_τ* + *o*(*τ*))^*n_k_* −1^. Since there are *n_k_* ways of this transition to occur, we have:

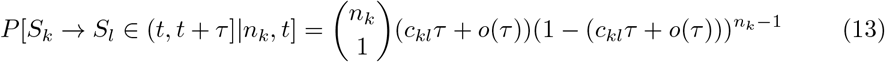

Eq 12 is then recovered as the *τ* terms of higher order are collected as *o*(*τ*) after expanding the binomial term in Eq 13.

**S2 Master equation** In this section, the steps for deriving Eq 5 are outlined. In principle, the master equation is based on the idea of mass conservation. First, the probability of observing a particular state-configuration at time *t* + *τ* is a function of the probabilities of the adjacent state-configurations at time *t* and the transitions between the corresponding probabilities of the state-configurations occurring in time step *τ*. The adjacent state configurations are defined as state-configurations with differences of +1 and *−*1 in two of the state counts, compared to the state-configuration of interest. The transitions between these corresponding probabilities are governed by the propensity function (*v*). Mathematically, the first step translates to:

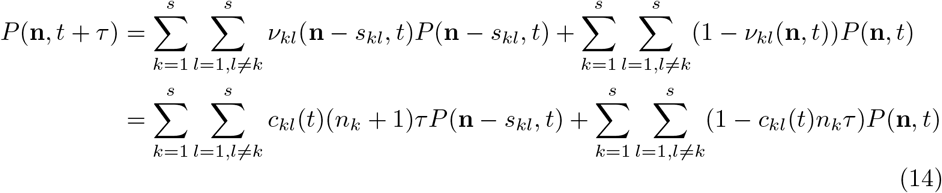

Rearranging Eq 14 and dividing it by *τ* yields:

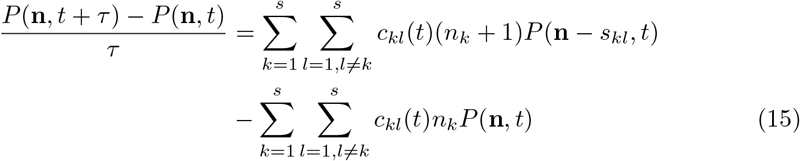

The master equation is then established after taking the limit *τ* → 0.

**S3 Generating function method** A probability generating function (PGF) for a vector **x** = (*x*_1_ *x*_2_ … *x_s_*) is defined by: 
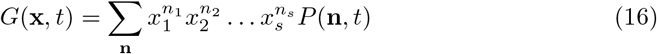

where 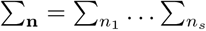. Differentiating *G*(**x**, *t*) with respect to *t* and assuming that the series is uniformly convergent, we obtain: 
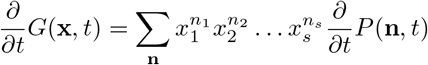
or 
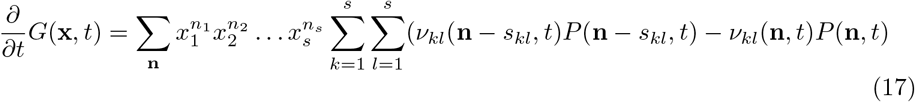
Eq 17 can be simplified by (1) recognizing the following identity (e.g., for *x*_1_): 
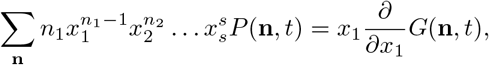
(2) using the definition of PGF (Eq 16), and (3) rearranging the summation index to obtain the following first-order linear partial differential equation (PDE): 
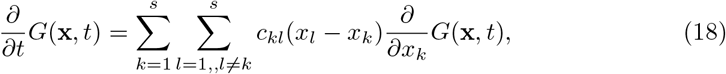
with an initial condition: 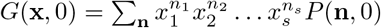 The PDE by solved by using the method of characteristics [14, 32]. The characteristics equations are: 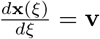 where **v** = (*v*_1_*v*_2_ … *v_s_*) and 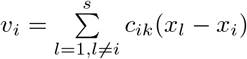. Putting *β*(*s*) = *G*(*x*(*ξ*), *t* − *ξ*), we have: 
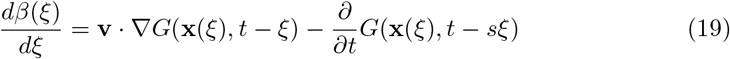
from which a general solution can be deduced: *G*(**x**(0), *t*) = *G*(**x**(*t*), 0) = *g*(**x**(*t*)). The system of the characteristics equations can be written as: 
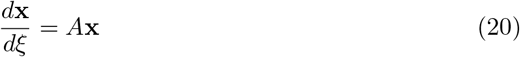

where *A* is an *s* × *s*, (*c_kl_*)_*k,l*∈{1,2,…,*s*}_, matrix with elements of the form: *A_kl_* = *c_kl_* − *γ_k_δ_kl_* with 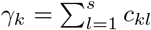 and *δ_kl_* is the usual Kronecker delta. Therefore, the solution of 20 takes the form: 
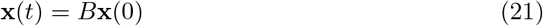
where 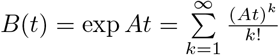, i.e. the matrix exponential. Given, *G*(**x**, 0), *G*(**x**(0), *t*) = *g*(*B***x**(0)). The solution of the PGF is then: 
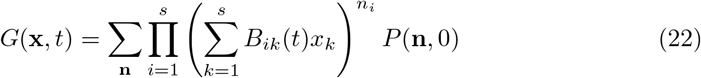

If we assumed all *n*_0_ individuals start in state *S*_1_, i.e. 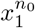, the solution of the PGF is then: 
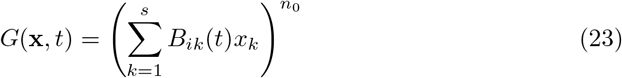

The probability density function of the state-configuration can be recovered from Eq 23 by using the definition of the PGF. We introduce a vector: **z**_*i*_ = [*z_i_*_1_ *z_i_*_2_ … *z_is_*] where *i* ∈ {1, 2, …, *s*} and the norm ∥**z**_*i*_∥ = *z_i_*_1_ + *z_i_*_2_ + … + *z_is_*. Using the multinomial theorem: 
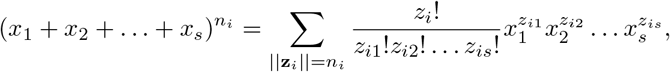
we write 
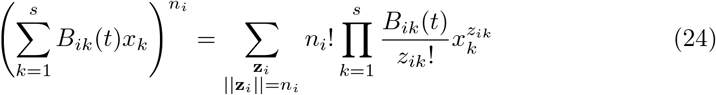

Therefore, we have: 
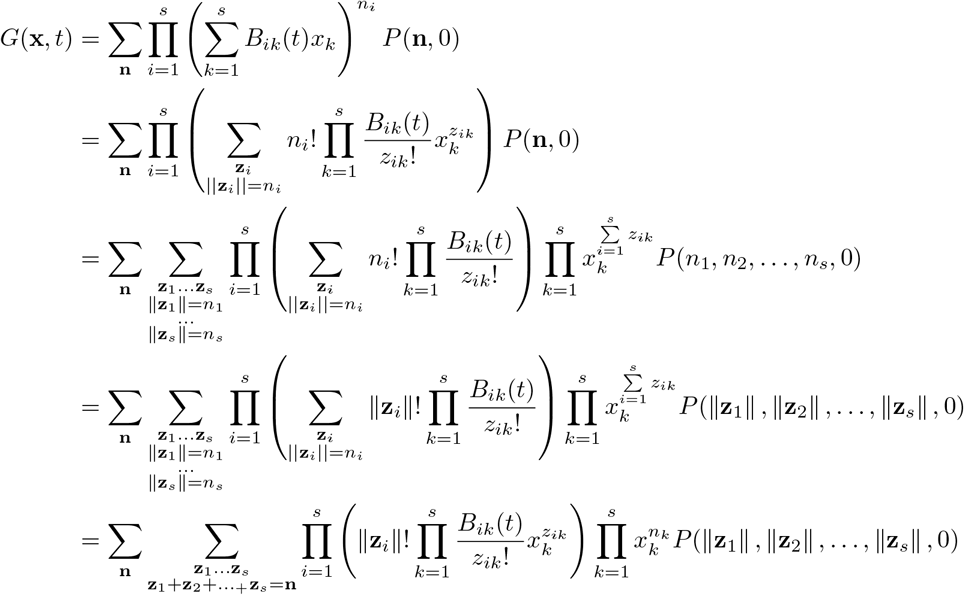

If we rearrange the last line, we have: 
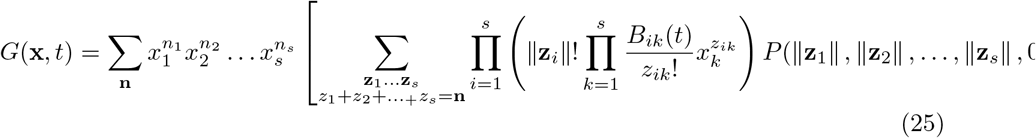

Therefore the solution to the master equation is obtained after equating the coefficients in Eq 16 with Eq 22: 
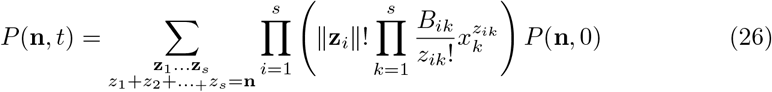

If we assumed all *n*_0_ individuals start in state *S*_1_, i.e. 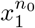, the solution takes the form of a multinomial distribution: 
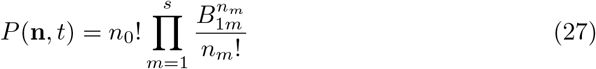

**S4 Mean and variance of the master equation** The first moment of the master equation can be computed by differentiating Eq 23 and set all *x* equal to 1, e.g. the mean of the number of individuals in state *S_i_*, given all *n*_0_ individuals start in state *S*_1_, is equal to: 
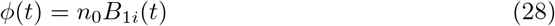

For an arbitrary initial distribution, the mean of the number of individuals in state *S_i_* is given by: 
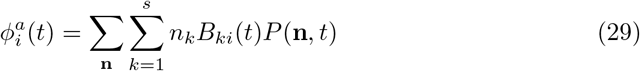

The variance of the master equation can be derived by using the following relationship: 
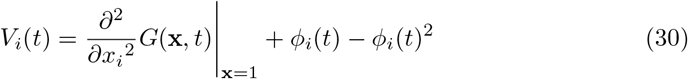

The variance of the number of individuals in state *S_i_*, given all *n*_0_ individuals start in state *S*_1_, is equal to: 
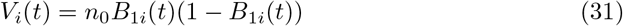

The variance of the number of individuals in state *S_i_* for an arbitrary initial distribution is given by: 
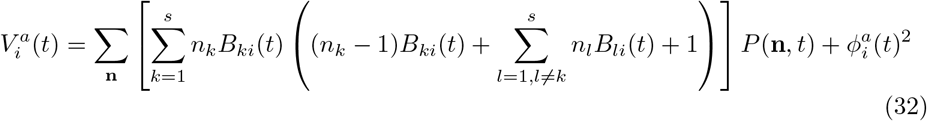

## Acknowledgments

I would like to thank Fernando Alarid-Escudero, Thomas A. Trikalinos, and Shaun P. Forbes for their insightful comments on the final draft.

